# Systems Analysis of de novo Mutations in Congenital Heart Diseases Identified a Molecular Network in Hypoplastic Left Heart Syndrome

**DOI:** 10.1101/2022.06.23.496464

**Authors:** Yuejun Jessie Wang, Xicheng Zhang, Chi Keung Lam, Hongchao Guo, Cheng Wang, Sai Zhang, Joseph C. Wu, Michael Snyder, Jingjing Li

## Abstract

Congenital heart diseases (CHD) are a class of birth defects affecting ∼1% of live births. These conditions are hallmarked by extreme genetic heterogeneity, and therefore, despite a strong genetic component, only a very handful of at-risk loci in CHD have been identified. We herein introduced systems analyses to uncover the hidden organization on biological networks of genomic mutations in CHD, and leveraged network analysis techniques to integrate the human interactome, large-scale patient exomes, the fetal heart spatial transcriptomes, and single-cell transcriptomes of clinical samples. We identified a highly connected network in CHD where most of the member proteins had previously uncharacterized functions in regulating fetal heart development. While genes on the network displayed strong enrichment for heart-specific functions, a sub-group, active specifically at early developmental stages, also regulates fetal brain development, thereby providing mechanistic insights into the clinical comorbidities between CHD and neurodevelopmental conditions. At a small scale, we experimentally verified previously uncharacterized cardiac functions of several novel proteins employing cellular assays and gene editing techniques. At a global scale, our study revealed developmental dynamics of the identified CHD network and observed the strongest enrichment for pathogenic mutations in the network specific to hypoplastic left heart syndrome (HLHS). Our single-cell transcriptome analysis further identified pervasive dysregulation of the network in cardiac endothelial cells and the conduction system in the HLHS left ventricle. Taken together, our systems analyses identified novel factors in CHD, revealed key molecular mechanisms in HLHS, and provides a generalizable framework readily applicable to studying many other complex diseases.

## Introduction

Congenital heart diseases (CHD), broadly defined by the structural and functional abnormalities in fetal heart, are the most common forms of birth defects and affects ∼1% of live births^1, 2^. Although CHD has a strong genetic component^3, 4^, the underlying genetic basis has largely remained elusive. Like many other pediatric diseases, large-scale copy number variations (together with aneuploidies) potentially explain ∼20% of CHD cases^4, 5^, and cases with monogenic causes could be solved by familial analyses^6, 7^. However, the genetic basis of sporadic cases, accounting for the majority of CHD probands, has largely remained unclear^3, 8^. The Pediatric Cardiac Genomics Consortium (PCGC) aims to fill in the knowledge gap by performing whole exome/genome sequencing on large-scale patient samples representing major sporadic CHD sub-types^9^. The latest PCGC study analyzed de novo and rare variants in the whole-exome data from 2,871 CHD probands and identified seven genes achieving genome-wide significance, together with a handful of genes showing suggestive associations with CHD^10^. These analyses collectively explained ∼10% of CHD cases in the cohort^10^. Targeting common variants, the latest genome-wide association analysis only identified one SNP reaching genome-wide significance^11^. Given the strong genetic basis in CHD, its complete genetic architecture has been yet to be discovered.

The existing analytical frameworks have been largely based on mutational recurrence analysis, where genes recurrently affected in probands than controls were identified for disease associations. However, in real clinical situations, different patients usually carry different sets of clinical mutations, and genes are more often individually than recurrently affected in patient populations. Importantly, these seemingly heterogeneous mutations usually functionally conserved onto common molecular pathways, giving rise to similar clinical phenotypes^12–15^. This is particularly the case in CHD, where its risk factors were more likely to affect different components in a shared molecular network, as opposed to recurrently affecting the same genes among patient populations^16, 17^. As such, it is not individual genes or mutations, but their structural organization on biological networks that defines the complete genetic architecture of the disease.

Our recent work has proposed a series of theoretic models to dissect convergent pathways in complex diseases from biological networks^12^. We herein leverage this system thinking to study CHD genomes. We developed a new framework to integrate network biology, genome analysis, spatial transcriptomics, and single-cell analysis for a direct revelation of the genetic basis in CHD. Combining computational and experimental approaches, we identified a highly connected cluster from the global human protein interaction network that was strongly implicated in fetal heart development and displayed significant enrichment for pathogenic mutations in CHD probands. Analyzing different CHD sub-types, the identified network was strongly associated with the hypoplastic left heart syndrome (HLHS) and displayed substantial dysregulation in cardiac endothelial cells and the conduction system in the under-developed left ventricle of the HLHS heart. We particularly note that the majority of the newly identified genes in this study had previously uncharacterized cardiac functions, nor their implications in CHD. Overall, our work provides a new systems framework for CHD genome analysis and has significantly expanded our knowledge about the genetic architecture of CHD.

## Methods and Materials

### An overview of the genomic resources

We analyzed 2,990 de novo mutations in cases and 1,830 de novo mutations in controls. These mutations were identified from the previous whole-exome sequencing study encompassing 2,871^10^ probands and 1,789 control individuals^18^, where the control sibling subjects were unrelated individuals to the probands. Among all the de novo mutations, we considered 323 and 129 loss-of-function (LoF) mutations (i.e. stop gain and loss mutations, and frameshift indels) in cases and controls, respectively given their clear functional consequences compared with missense mutations. At the gene level, we considered those intolerant to LoF mutations with a pLI score^19^ greater than 0.8, and therefore the presence of de novo LoF mutations in these genes is a strong indicator of mutational pathogenicity. With this, we identified 120 and 35 genes in the proband and control cohorts, respectively (**Table S1**), for subsequent analyses. Gene functional enrichment test confirmed a significant overrepresentation of genes regulating heart development in the 120 proband genes but not in the control genes, and we therefore considered them as CHD candidate genes. Gene function enrichment analyses throughout this work were performed using Enrichr^20^ (Database: as of Oct, 2021).

### Analysis of the protein interaction network

We seeded the 120 proband proteins in the human protein interaction network. The network was analyzed in our previous publication^13^ with 16,085 unique proteins (or genes) and 217,605 interactions. We implemented the random walk algorithm (personalized page rank^21^) on the network by setting the restart probability at 0.1 and the maximum number of iterations at 500. For each node on the network, we derived its probabilities of visiting all other nodes on the network and a greater probability indicates greater reachability between two nodes, thereby increased topological distance. To identify those topologically closer to the 120 CHD candidate proteins seeded on the network, we performed Wilcoxon rank-sum test to determine whether a given node is more reachable to the 120 CHD candidate proteins relative to its reachability to all other proteins on the network. We performed the test on each of the nodes on the network (excluding the 120 CHD proteins), and the derived P values were corrected by Benjamini-Hochberg (false discovery rates less than 0.05). Therefore, on the proteome scale, we agnostically identified another set of 120 new proteins forming a highly connected network the 120 seeded PCGC proteins. To determine the identification of the new proteins was not expected by chance, we performed degree-preserving shuffling^22^ to permute the protein interaction network, and recorded the number of nodes in the largest connected component in each permutation simulation. The observation from the real dataset cannot be observed from the permutation test, thereby statistical significance suggesting biological functions.

### Gene expression analysis

We analyzed the time-course gene expression data in the mouse cardiogenesis process^23^, where human-mouse orthology mapping was based on the Ensembl Biomart annotation, and we only considered those with one-to-one orthology mapping. Expression of gene symbols mapped onto multiple probesets identifiers were averaged. Gene expression was then normalized across time points followed by a hierarchical clustering, revealing two expression clusters on the network, Group-I (G-I) and Group-II (G-II), in **Figure 2B**. We also analyzed gene expression in the developing fetal heart from postconceptional day 96 (gestational week 13.7) to 147 (gestational week 21) using microarray data from the NIH Roadmap Epigenomics Mapping Consortium^24^ (GSE18927 in GEO). All probesets intensities were normalized onto a logarithm scale, and signals from probesets mapped onto the same gene symbol were averaged. At each time point, we compared expression of genes of interest against the transcriptome background to determine their molecular activities and only protein coding genes were used in this comparison. To confirm our observation, we also performed analysis using RNA-seq data in fetal heart samples in postconceptional weeks 19 and 28^25^ (ENCSR000AEZ from the ENCODE consortium: https://www.encodeproject.org). Statistical comparisons were determined by the Wilcoxon rank-sum test.

### Spatial transcriptome analysis

We analyzed the spatial transcriptome data in the fetal heart from a recent publication^26^, and analyzed spatial RNA-seq data in 3,115 tissue spots (across sections) at 4.5-5, 6.5 and 9 postconceptional weeks (PCW). We standardized gene expression on spots from all three postconceptional weeks by NormalizeData (settings: normalization.method = “LogNormalize”, scale.factor = 10000) from Seurat package^27^ (version 4.0.4). We then performed quantile normalization across all tissue spots across all sections. In each tissue section, the original study clustered tissue spots into groups with shared transcriptome profiles, which corresponded to different anatomical compartments in the developing heart. To quantify region-specific gene expression in the heart, we averaged expression of each gene across spots within the same anatomical compartments of a fetal heart across all sections. To determine statistical significance, we compared genes of interest against the transcriptome backgrounds (protein-coding genes) in each annotated anatomical compartments using Wilcoxon rank-sum test. For visualization, we used one representative tissue section with the most spots at each PCW.

### Whole-exome analysis of PCGC proband cohort

To determine the enrichment of pathogenic mutations on this network genes among CHD proteins, we examined the PCGC whole-exome-sequencing data in dbGAP (as of Feb, 2021, dbGAP-24034, gap_accession: phs000571, gap_parent_phs: phs001194, SRP025159). We used the same control subjects as described in the original study^10, 18^, and downloaded the whole exome sequencing data from SFARI BASE (https://www.sfari.org/resource/sfari-base/). The analyzed individual identifiers were separately listed in **Table S6**. We downloaded the raw FastQ data files from dbGAP, and performed variant calls following the Best Practice procedure recommended by GATK. We utilized the Sentieon toolkit which substantially increases the computational efficiency while keeping genotyping accuracy^28^. We performed independent quality control analyses to ensure high quality of the called variants. We utilized VCFtools (http://vcftools.sourceforge.net) to compute the distribution of Ti/Tv ratios across all analyzed individuals (shown in **Figure S5**).

Using all the called exonic variants, we performed principal component analysis (PCA) by aggregating individuals from case and control cohorts. The PCA analysis ensured almost identical population structure between cases and controls. For rare variants, we only considered those absent from the 1000 Genome Database (https://www.internationalgenome.org). We elected to use 1000 Genome as our reference, instead of the Genome Aggregation Database (gnomAD^29^) or TOPMed^30^, because a significant portion of samples in gnomAD and TOPMed were individuals with cardiovascular diseases, such as the those in Atrial Fibrillation Genetics Consortium (AFGen) and Myocardial Infarction Genetics Consortium. We annotated the called variants using the ANNOVAR package^31^, which provided CADD (Combined Annotation Dependent Depletion) phased scores (Database: CADD v1.6 as of Feb, 2021) for each of the called variants^32, 33^. For each personal exome in this study (in case and control cohorts), we computed the mean CADD scores for non-synonymous (LoF and missense) variants affecting the network genes, and then compared the mean CADD score distribution among all probands in each CHD subtype against individuals in the control cohort. The same comparison was also performed on synonymous variants as a set of negative controls. As another set of negative control, we obtained 62 lung-specific protein-coding genes from a previous publication^34^. To determine whether the observed mutational pathogenicity was specific to Group-II genes, we performed a permutation analysis, where, in each permutation, we randomly sampled rare non-synonymous variants from the exome background in the HLHS cohort, matching the number of rare non-synonymous variants in Group-II genes in the same HLHS cohort. We performed the same sampling on the control cohort, and then compared CADD scores between the two randomly sampled variant list. We performed the permutation 100 times, and confirmed that CADD scores were not statistically significant between the randomly sampled variant sets from case and control cohorts, respectively (p = 0.98, permutation test).

### Analysis of single-cell RNA-seq data from an HLHS heart

We re-analyzed the published single-cell RNA-seq data in an HLHS fetal heart^35^, and specifically performed our comparison on the under-developed left ventricle against that of the matched control sample. We followed the cell type clustering described in the original publication^35^. For each gene, we averaged its expression across all cells in a given cell type, and for each cell type, we asked whether the network genes (Group-I and II genes) tended to have increased expression relative to the transcriptome background. Wilcoxson rank-sum test was used to determine statistical significance.

### iPSC conversion to cardiomyocytes

All induced pluripotent stem cell (iPSC) lines were reprogrammed by the Sendai virus expressing 4 Yamanaka factors (CytoTune®-iPS Sendai Reprogramming Kit, Invitrogen) as previously described^36^. Protocols in the study were approved by Human Subjects Research Institutional Review Board (IRB) at Stanford University. Human iPSCs were maintained in Essential 8™ Medium (Gibco®, Life Technology). For passaging, iPSC culture was dissociated with Accutase (Innovative Cell Technologies) at 37°C for 15-20 min, and suspended iPSCs were reseeded on Matrigel-coated plates (BD Biosciences, San Jose, CA) at a density of 500 K cells per well in 6-well plates.

Beating induced pluripotent stem cell-derived cardiomyocytes (iPSC-CMs) were generated using the RPMI + B27 method as described^37, 38^. Briefly, human iPSCs were kept in 6-well plates until passage 20. Once they reached > 80% confluence, the medium was switched to RPMI 1640 with Insulin minus B27 supplement (Gibco®, Life Technology) and 6 µM CHIR99021 (Selleckchem) for 2 days. They were allowed for 1-day recovery with RPMI + B27 (minus insulin) medium. Cells were then treated with 5 µM IWR-1 (Sigma) for 2 days, and then fresh RPMI + B27 (minus insulin) medium for another 2 days. Cells were then switched to RPMI + B27 medium for 3 days. Beating cardiomyocytes were purified for 2-3 rounds of 3-day glucose-free RPMI + B27 medium treatment.

To downregulate the expression of our target genes, we performed siRNA transfection in iPSC-CMs as described previously^39^. 80 μl of siRNA (Dharmacon) was added into a master mix of 3.2 μl DharmaFECT1 (Dharmacon) transfection reagent and 236.8 μl OptiMEM (Thermo Fisher Scientific) and incubated for 20 min before addition to a 6-well plate of iPSC-CMs. The cell media was then changed after 24 hr. Cells were then subjected to contractility assays or harvested 48 hr after medium change.

### Cellular contractility assays

To assess iPSC-CM contractility, iPSC-CMs were re-seeded on Matrigel-coated 24-well plates and cultured for 7 days to recover their spontaneous beating, as previously described^40^. Contraction of monolayer cardiomyocytes was recorded with high resolution motion capture tracking using the SI8000 Live Cell Motion Imaging System (Sony Corporation). During data collection, cells were maintained under controlled humidified conditions at 37°C with 5% CO2 and 95% air in a stage-top microscope incubator (Tokai Hit). Functional parameters were assessed from the averaged contraction-relaxation waveforms from 10-sec recordings.

### Western blotting

Human iPSCs and iPSC-CMs grown in 6-wells plates were harvest and lysated in RIPA buffer with Complete Mini, EDTA-free Protease inhibitor cocktail tablets (Roche). The lysates were placed on ice for 30 minutes, followed by centrifuging at 14000 rpm for 20 min, the supernatants were then collected as proteins. BCA Protein Assay kit (Thermo Fisher Scientific) was used to measure the protein concentration. Western blot was performed according to the standard protocol. Briefly, Equal amounts of protein was treated by SDS-PAGE electrophoresis and transferred to a nitrocellulose membrane. After nonfat milk blocking, the membrane was incubated with the following primary antibodies at 4°C overnight, respectively: TLK1 (Cell Signaling Technology, 4125S; 1:1000 dilution), TEAD2 (Proteintech, 21159-1-AP; 1:500 dilution), RBBP5 (Bethyl Laboratories, A300-109A-M; 1:1000 dilution), ASH2L (Bethyl Laboratories, A300-107A-M; 1:1000 dilution) and GAPDH (Abcam, ab8245; 1:1000 dilution). Subsequently, the membrane was incubated in protein-specific HRP conjugated secondary antibody for 1 hr at room temperature. Restore Western Blot Stripping Buffer (Thermo Fisher Scientific) was used to clean RBBP5 antibody for detecting ASH2L protein. The blots were visualized using chemiluminescence (Thermo Fisher Scientific).

### Cell sorting by FACS

Flourescent activated cell sorting (FACS) was performed to determine the cardiomyocyte differentiation efficiency. iPSC-CMs were dissociated and stained with cardiac troponin T antibody (ab45932, Abcam), and goat anti-Rabbit secondary antibody (A32731, Thermo Scientific) using Fixation/Permeabilization Kit (554714, BD Biosciences). The stained cells were filtered through a 35 µm cell strainer snap cap and collected in a 5 ml FACS tube (Corning). The cells were analyzed on a BD Biosciences FACS Aria II instrument using FACSDiva software. The cells were gated on the basis of forward scatter and side scatter. Flow cytometric gates were set using parental cells.

### siRNA experiments and RNA-seq

To generate ASH2L knockout iPSCs, CRISPR/Cas9 gene editing was performed using two single-guide RNA (sgRNAs) flanking the exon 4-5 region of ASH2L. The guide DNA oligos were designed using a web-based tool (crispr.mit.edu/) and chosen based on a high score for on-target binding and the lowest off-target score. The gRNAs were cloned into the pSpCas9(BB)-2A-GFP vector (PX458; a gift from Feng Zhang; Addgene plasmid #48138) using annealed reverse complementary guide DNA oligos. The sequences of the sgRNAs were gRNA_3’: GGCAGAGACGGATGCAACAG and gRNA_3’: GTGGTTGTATAACAGAATAT. Two CRISPR/Cas9 vectors (1 ug each) were transfected in hiPSCs (SCVI480 using the Lipofectamine Stem Transfection Reagent (Thermo Fisher Scientific). The cells were dissociated using TrypLE express 1x (Thermo Fisher Scientific) and GFP+ cells were sorted by flow cytometry 24 hr post-transfection. GFP+ cells were seeded at a density of 1,000 cells per well in a 6-well plate to generate clonal isolates. Ten to fourteen days after seeding, 48 individual clones were picked for genotypic screening by PCR. (FW: AGCTAGTTTCAGAGTCCAAGATAAA, RV: GATGGAGAAAGAAGCTATAGAGGAG) Knock-out clones were confirmed by DNA agarose gel. Heterozygous clones F5 and H12 are selected for subsequent experiments.

For RNA-seq, NEBNext Ultra II RNA kit with PolyA Selection was used to construct RNA library. Two replicates were completed for each condition. Then standard RNA-seq protocols were followed and we generated > 28 million 76 bp single end reads per sample. The original reads were mapped to human genome (GRCh37) with STAR 2.7.3a^41^ with default settings, then reads with MAPQ > 20 were collected for further analysis. FeatureCounts^42^ was applied to count the number of reads mapping to genomic feature. Differentially expression was detected by DESeq2^43^. GO terms were enriched with clusterProfiler^44^.

## Results

### NetWalker identified a sub-network in CHD

We analyzed de novo mutations identified from the PCGC whole-exome sequencing data from 2,871 CHD probands^10^. We agnostically identified 120 genes from the PCGC CHD probands that were affected by de novo loss-of-function (LoF) mutations (stop gain and loss mutations and frameshift indels), and LoF mutations in these genes were highly depleted in natural human populations (see **Methods and Materials** and **Table S1A**), suggesting their sensitivity to gene copy loss. The 120 genes displayed an overall functional enrichment for abnormal heart development (FDR = 4.7e-4), decreased fetal cardiomyocyte proliferation (FDR = 2.2e-3) and abnormal cardiovascular system morphology (FDR = 6.5e-3) (**Table S1B**), well representing CHD candidate genes. With exactly the same procedure, we also identified 35 genes from the accompanied control cohort from the original publication^10^, which, as expected, did not show enrichment for cardiac functions (**Table S1C** and **D**). We examined the topological occupancies of these identified PCGC candidate genes on the high-quality experimentally derived human protein interaction network that was compiled and quality-checked in our previous publication^13^, encompassing 16,085 unique proteins (or genes) and 217,605 interactions (see **Methods and Materials**, **Table S2**). We observed that these PCGC candidate proteins tended to have significantly increased connectivity compared with those identified from unaffected sibling controls (p = 0.02, Wilcoxon rank-sum test, **Figure 1A**), thereby occupying the central topological positions on the interaction network. We confirmed this observation using genes affected by de novo synonymous mutations in the PCGC probands and unaffected siblings, respectively: as expected, no significant difference was observed (p = 0.24, **Figure 1A**). This comparison suggested a global impact of the identified 120 PCGC proteins given their central topological positions on the network. We next asked whether these identified proteins were topological neighbors on the interaction network, which would help reveal convergent molecular pathways in CHD. We calculated the fraction of the identified 120 proteins that were direct interacting partners on the network, which indeed displayed a significant enrichment compared to the genes identified from the unaffected siblings using the same procedure (p = 6.8e-3, Fisher’s exact test, **Figure 1B**), and the pattern was absent from control genes affected by de novo synonymous mutations (p = 0.19, Fisher’s exact test, **Figure 1B**). Taken together, although the 120 genes were individually and agnostically identified from the PCGC CHD exomes, their topological positions on the global interaction network were not random: (1) they tended to occupy central positions on the network to exert their global impact on biological processes; (2) they were more likely to interact with each other on the biological network, suggesting their formation of a functional convergent sub-network underlying heart development.

**Figure 1.**
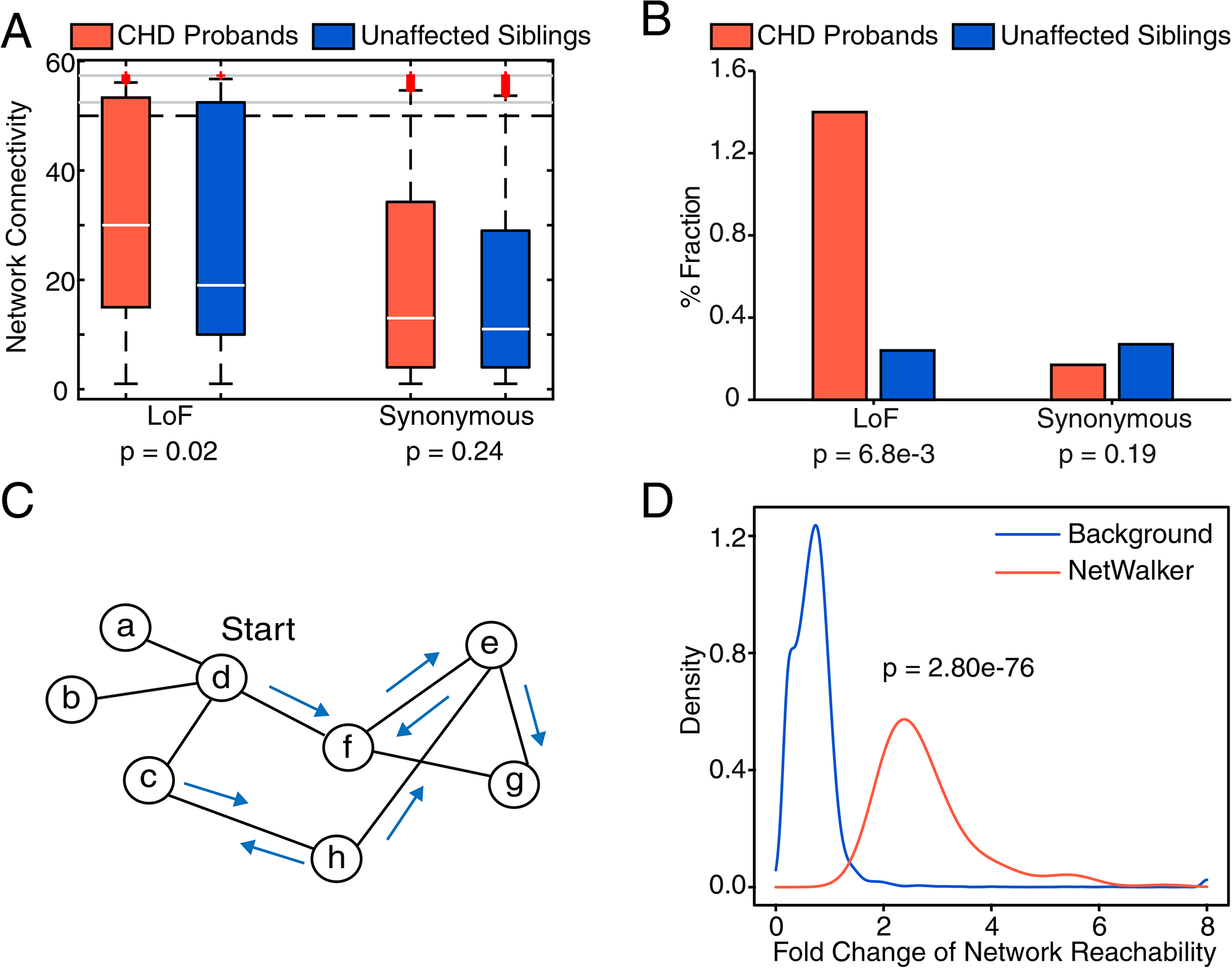
NetWalker Identified a highly connected network in CHD. **A.** PCGC candidate genes tended to occupy central positions on the protein interaction network. PCGC candidate genes (red) were identified by dosage-sensitive genes affected by de novo loss-of-function (LoF) mutations from the PCGC CHD cohorts. The same procedure also identified affected genes (blue) from the matched unaffected sibling cohort. Network connectivity indicates the number of interacting partners for each protein on the network. As an independent control experiment, the same comparison was also performed on genes affected by de novo synonymous mutations, whose functions were presumably neutral. P-values were derived from Wilcoxon rank-sum test. **B.** PCGC candidate genes were more likely to maintain mutual interactions on the network. The fractions of interacting proteins were computed among the candidate and control genes identified from proband and sibling cohorts, respectively. Genes affected by de novo synonymous mutations were used as an independent control experiment. P-values were derived from the Fisher’s exact test. **C.** The schematic presentation of the random walk algorithm on the protein interaction network. The random walk scheme starts from every node on the network following stochastic flow till convergence. For each protein, the probabilities of visiting all other proteins on the network will be calculated, which defines the reachability of this node to any other nodes on the network. **D.** The identified network component has substantially increased reachability to PCGC CHD candidate proteins relative to all other proteins on the network. P-values were derived from Wilcoxon rank-sum test.

We developed the NetWalker algorithm to dissect the complete structure of the underlying convergent molecular network in CHD seeded by the 120 candidate proteins (termed the PCGC proteins/genes). NetWalker is essentially a random walk scheme on a complex network which calculates the probability of visiting one node from any other nodes assuming stochastic flow on the network (the “reachability” between any two nodes on the network, **Figure 1C**)^45, 46^. Specifically, for each protein on the protein interaction network, we computed its reachability to each of the PCGC proteins, as well as to the entire collection of the human proteins on the network. At a false discovery rate of 0.05 and fold change of 2, we agnostically identified another set of 120 new proteins (the NetWalker proteins/genes) topologically more reachable to the 120 PCGC proteins than to the global proteome background (**Figure 1D**). Most of these 120 newly identified NetWalker proteins had not been reported by previous CHD mutational analyses^10, 47^, and we next sought to functionally characterize their potential functions in heart development.

These newly identified 120 proteins have extensive interactions with the candidate PCGC proteins, where 179 (88/120 from the PCGC proteins and 91/120 from the NetWalker proteins) out of the 240 proteins (120 PCGC+120 NetWalker) formed a highly connected network (**Figure 2A**). To demonstrate statistical significance of the identified network, we performed degree-preserving shuffling^22^ on the network, where we randomly permutated the interacting partners for each protein on the interaction network while maintaining their respective connectivity. With the same set of seed proteins, implementing NetWalker 100 times on these permuted networks only identified highly fragmented sub-networks (**Figure S1**), suggesting that the identified highly connected network (**Figure 2A**) was not expected by chance. Excluding the seed PCGC candidate proteins, throughout our analyses we sought to characterize the newly identified NetWalker proteins (the orange nodes in **Figure 2A**) for their biological significance in regulating heart development.

**Figure 2.**
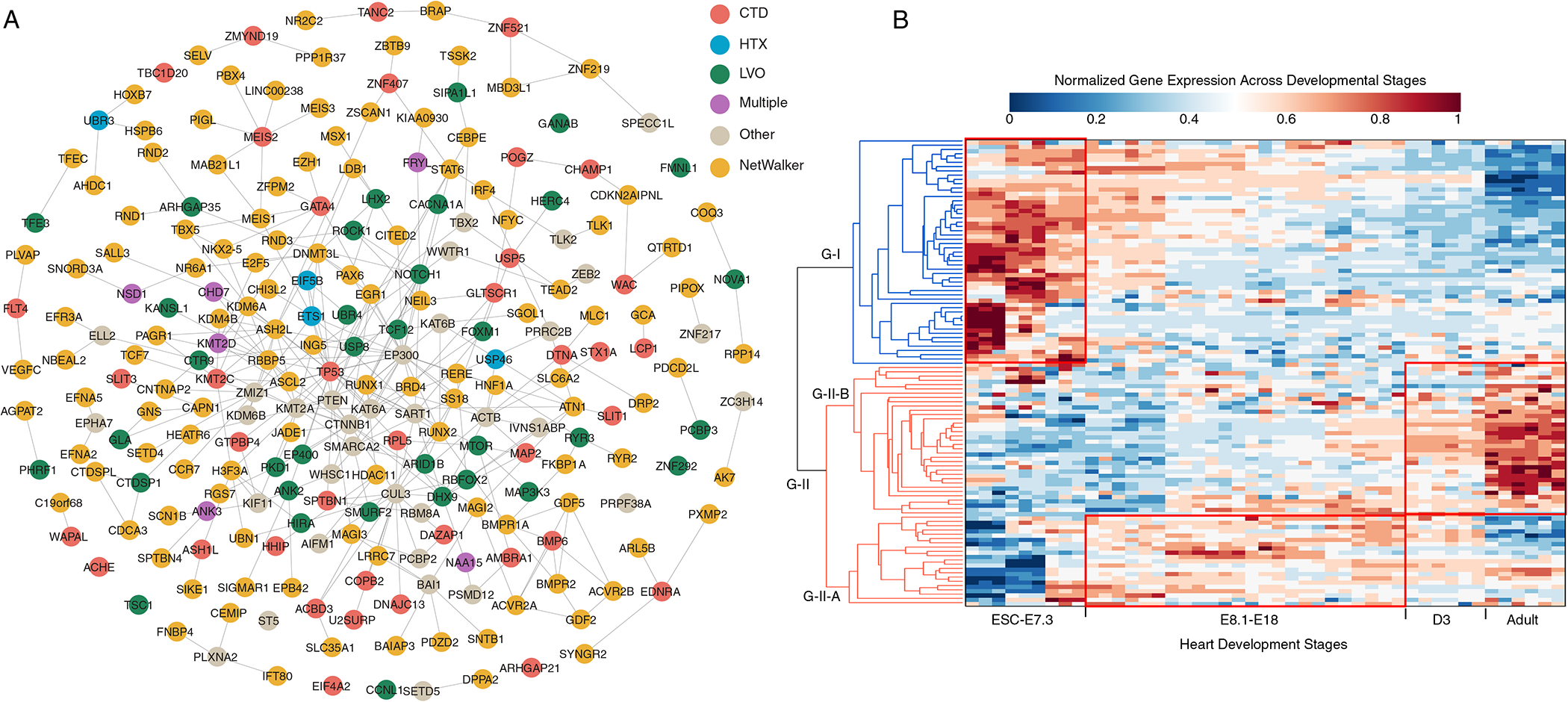
Functional characterization of the CHD network. **A.** An overview of the identified network seeded with PCGC candidate proteins (grouped by CHD subtypes that were color coded), and the orange nodes were novel proteins identified by the NetWalker algorithm. The subtype annotations were derived from the original publication^10^, where CTD, HTX and LVO stand for conotruncal defects, heterotaxy and left ventricular outflow tract obstruction, respectively. Other indicates more than one subtype was associated with the corresponding proteins. **B.** Temporal expression of the network genes across heart developmental stages. Human genes were mapped onto mouse orthologs, and hierarchical clustering revealed two expression components of the network, where Group-I (G-I) genes displayed preferential expression from embryonic stem cells (ESC) to E7.3, whereas Group-II (G-II) genes exhibited substantial expression enrichment from E7.3 to postnatal and adult stages. Close examination of Group-II genes further revealed two subcluster structure, where Group-II-A (G-II-A) was preferentially expressed across fetal developmental stages and Group-II-B (G-II-B) was more specific in postnatal stages, particularly strong in the adult heart.

### Functional characterization of the CHD network

Although the newly identified NetWalker proteins had no significant mutations from previous mutational analyses in the PCGC cohort (the orange nodes in **Figure 2A**), several known CHD risk factors (most from previous clinical studies) could be immediately recognized, including TBX5^48^, NKX2-5^49^, CITED2^50^, IFT80^51^, ZFPM2^52, 53^ and ACVR2B^54^. Additionally, our algorithm also identified MSX1, whose association with CHD was recently suggested by a CHD GWAS study^11^. Despite these known proteins, to the best of our knowledge, the majority of the proteins identified by NetWalker (**Figure 2A**) had uncharacterized function in heart development nor in CHD. We investigated their gene expression in the developing heart using mouse models and the human fetal heart. We first considered the time-course transcriptomic data during cardiogenesis in mouse, where transcriptomes in the developing heart were densely sampled and profiled across key heart developmental stages, from embryonic stem cells to fetal, postnatal and adult stages^23^. Utilizing one-to-one unambiguous mouse-human orthology mapping, we observed that the NetWalker genes formed two expression clusters across the heart developmental stages: Group-I (G-I) genes displayed preferential expression from embryonic stem cells (ESCs) to E7.3, whereas Group-II (G-II) genes exhibited substantial expression enrichment from E7.3 to postnatal and adult stages (**Figure 2B**). Note that the heart tube forms at ∼E7.3 (corresponding to ∼2.5 postconceptional weeks, PCW, in humans). Close examination of Group-II genes further revealed two subcluster structure, where Group-II-A (G-II-A) was preferentially expressed across fetal developmental stages and Group-II-B (G-II-B) was more specific in postnatal stages, particularly strong in the adult heart (**Figure 2B**). It is also important to note that the observed expression propensities were relative comparisons across developmental stages, and therefore the increased expression in particular time points do not necessarily preclude its molecular activities in other time points.

We further leveraged the recently published spatial transcriptomic data in the fetal heart^26^ to investigate the molecular activities of the identified genes in specific heart compartments. The original experiments evenly sampled tissue spots (each containing ∼30 cells) from the fetal heart of serial sections in postconceptional weeks (PCW) 4.5-5, 6.5 and 9, and determined the transcriptome in each tissue spot so as to gain insights into the spatial effects on modulating gene expression during heart development^26^. These tissue spots were then clustered by their transcriptome similarity to reveal cell architecture defining spatial compartments of the developing fetal heart. We analyzed our identified genes in these spatial compartments, and did not observe significant expression enrichment for Group-I genes across all heart compartments in PCW 4.5-5, 6.5 and 9. The observation was expected given their early embryonic expression in the mouse data (**Figure 2B** before ∼E7.3). The lack of expression enrichment was also observed for Group-II-B genes (**Figure 2B**), consistent with their postnatal and adult expression in the mouse data. However, the Group-II-A genes displayed strong expression enrichment across all fetal heart compartments in both PCW 6.5 and 9 (**Figure 3A and B**), but not in PCW 4.5-5. This again demonstrated the overall concordance with their fetal expression in the developing mouse heart (**Figure 2B**), but more precisely suggested their developing timing after PCW 4.5-5. Overall, these spatial transcriptome data supported our observation from the mouse data.

**Figure 3.**
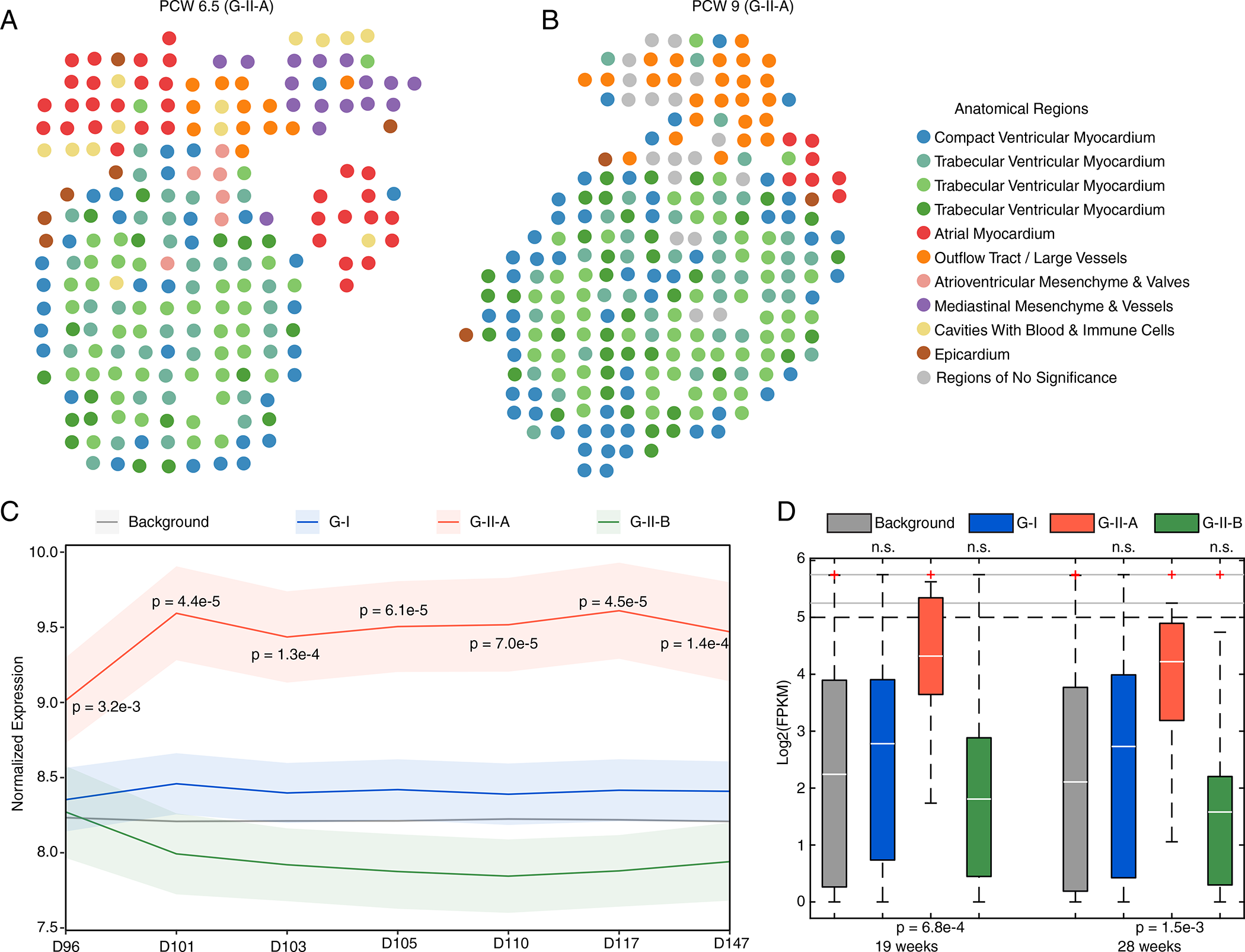
Spatiotemporal expression analysis of the identified CHD network. **A-B.** Group-I (G-I) genes and Group-II-B (G-II-B) genes did not show significance across all time points. Group-II-A (G-II-A) genes showed pervasive expression activities across most spatial spots in the fetal heart in both PCW 6.5 (**A**) and 9 (**B**). **C**. Group-II-A (G-II-A) genes showed significantly increased expression in the fetal heart from postconceptional day 96 to day 147, whereas Group-I (G-I) genes and Group-II-B (G-II-B) genes did not display statistical significance (p > 0.05, Wilcoxon rank-sum test). **D**. Group-II-A (G-II-A) genes showed significantly increased expression in the fetal heart in the gestational weeks 19 and 28 based on RNA-seq data. The statistical significance was not observed from other gene groups.

Leveraging resources in the Epigenome Roadmap Project^24^, we further analyzed the fetal heart transcriptome data in 7 discrete time points from postconceptional day 96 (gestational week 13.7) to day 147 (gestational week 21), and again we observed strong molecular activities of the Group-II-A genes across all time points relative to the transcriptome background (**Figure 3C**). The significance was absent for Group-I and Group-II-B genes. We further examined RNA-seq data from the ENCODE project^25^ for the fetal heart in gestational weeks 19 and 28, again confirmed strong activities of the Group-II-A genes in both gestational weeks (**Figure 3D**). Taken together, all the human data confirmed molecular dynamics of the identified genes as we observed from the mouse data: Group-I genes were specific to early embryonic stages; Group-II-A genes were specific for fetal development, whereas Group-II-B genes were more specific for the postnatal and adult heart.

We performed gene ontology analysis on the 120 NetWalker proteins (the orange nodes in **Figure 2A**) to determine their molecular functions in modulating heart development. Consistent with the above observation on gene expression, Group-II genes displayed significant functional enrichment for outflow tract septum morphogenesis, ventricular septum development and regulation of cardiac muscle cell proliferation (**Table 1**). The enrichment for heart functions was also observed when splitting Group-II genes into two sub-groups (Group-II-A and Group-II-B). For Group-I genes, their functional enrichment was significant for heart development, particularly in right atrial isomerism and abnormal cardiovascular development; however, unexpectedly, their gene ontology also displayed unexpected enrichment for neural functions, especially for open neural tube defects and exencephaly (**Table 1**). Given the significant comorbidities between CHD and neurodevelopmental diseases^55, 56^, particularly with neural tube malformations (e.g. 40.6% of probands with spina bifida aperta develop CHD^57^), this observation likely suggested the underlying etiological causes (which will be experimentally verified below). Because Group-I genes were specific to early developmental stages and the enrichment for brain functions was completely absent from Group-II genes (**Table 1**), the shared molecular etiologies between CHD and neurodevelopmental disorders should occur only at the early developmental stages.

**Table 1.**
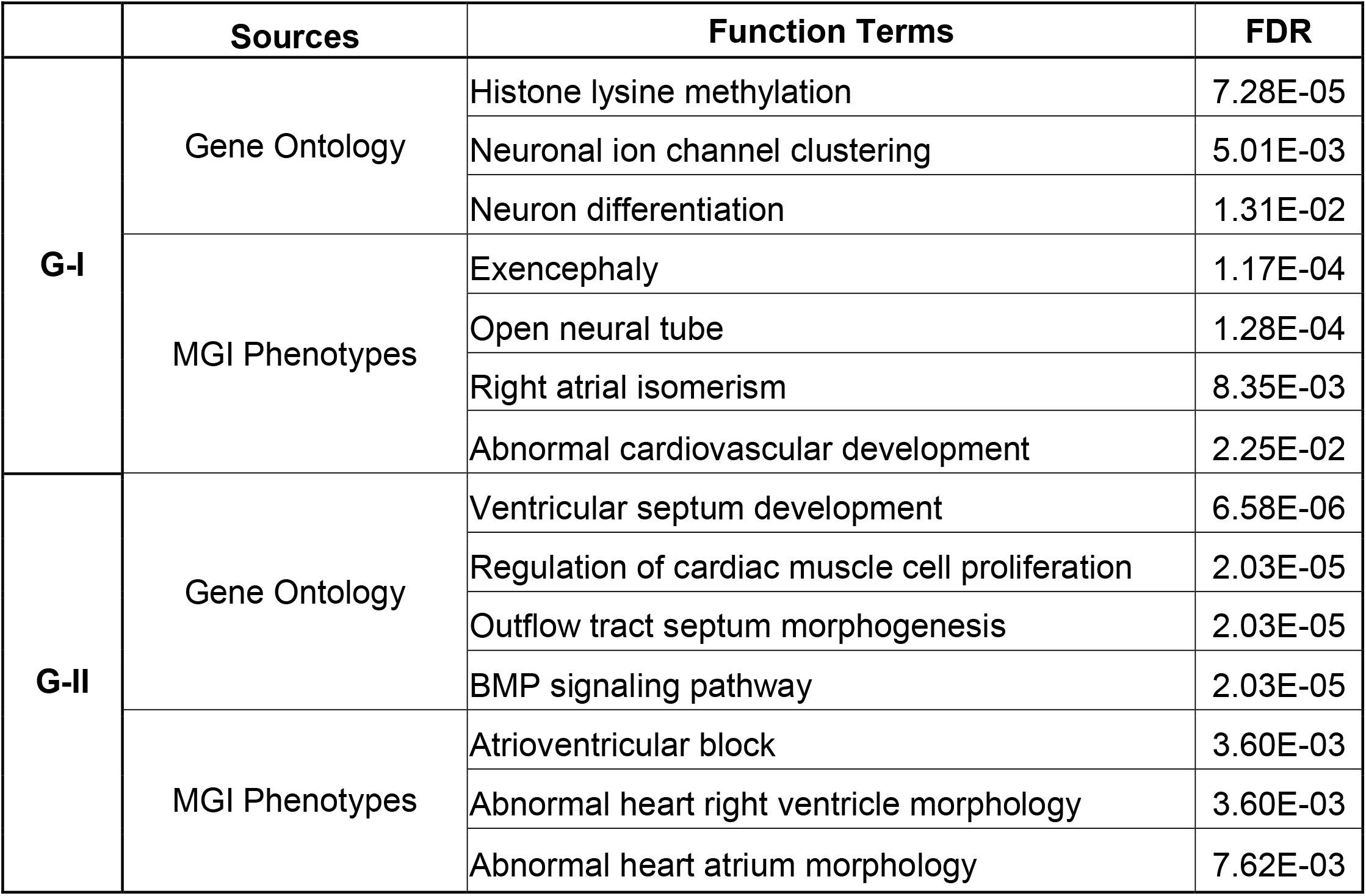
Enriched Gene Ontology Terms for Group-I and II Genes on the Network.

|  | Sources | Function Terms | FDR |
| --- | --- | --- | --- |
| **G-I** | Gene Ontology | Histone lysine methylation | 7.28E-05 |
|  |  | Neuronal ion channel clustering | 5.01E-03 |
|  |  | Neuron differentiation | 1.31E-02 |
|  | MGI Phenotypes | Exencephaly | 1.17E-04 |
|  |  | Open neural tube | 1.28E-04 |
|  |  | Right atrial isomerism | 8.35E-03 |
|  |  | Abnormal cardiovascular development | 2.25E-02 |
| **G-II** | Gene Ontology | Ventricular septum development | 6.58E-06 |
|  |  | Regulation of cardiac muscle cell proliferation | 2.03E-05 |
|  |  | Outflow tract septum morphogenesis | 2.03E-05 |
|  |  | BMP signaling pathway | 2.03E-05 |
|  | MGI Phenotypes | Atrioventricular block | 3.60E-03 |
|  |  | Abnormal heart right ventricle morphology | 3.60E-03 |
|  |  | Abnormal heart atrium morphology | 7.62E-03 |

### Experimental validation confirmed the role of the novel proteins in heart development

Given the clear and strong implication of Group-II genes in heart development (**Figure 3** and **Table 1**), we next closely examined how Group-I genes modulate heart development, particularly those also implicated in neurodevelopmental disorders. We followed our previous practice^15^ and partitioned the identified network (**Figure 2A**) into 33 non-overlapping topological clusters (**Table S3**), where in each cluster, proteins were densely connected with their interacting partners but sparsely interacted with proteins outside of their respective clusters. This approach has enabled us to understand each protein’s function in the context of its own interaction module. Cluster #4 is an excellent example to demonstrate the convergence of CHD-associated mutations (**Figure 4A**), where 12 out of the 15 member proteins were affected by de novo LoF mutations identified from CHD proband (the PCGC candidate genes). These genes were heralded by FOXM1, whose mouse mutants displayed ventricular hypoplasia, cardiomyocyte deficiencies^58^ and many other cardiac anomalies^59^. Although these de novo mutations were individually identified from different CHD probands, their topological clustering in the same module demonstrated the mutational convergence in CHD onto a common component. As such, for the remaining three novel proteins in the same cluster (SLC6A2, DRP2 and MLC1, **Figure 4A**) identified by our NetWalker algorithm, it is reasonable to expect their function in modulating heart development. Indeed, SLC6A2 mouse mutants display smaller heart sizes with increased heart rate (Mouse Genome Informatics^60^), and DRP2 is specific to heart-derived endothelial cells among many other endothelial cell types^61^.

**Figure 4.**
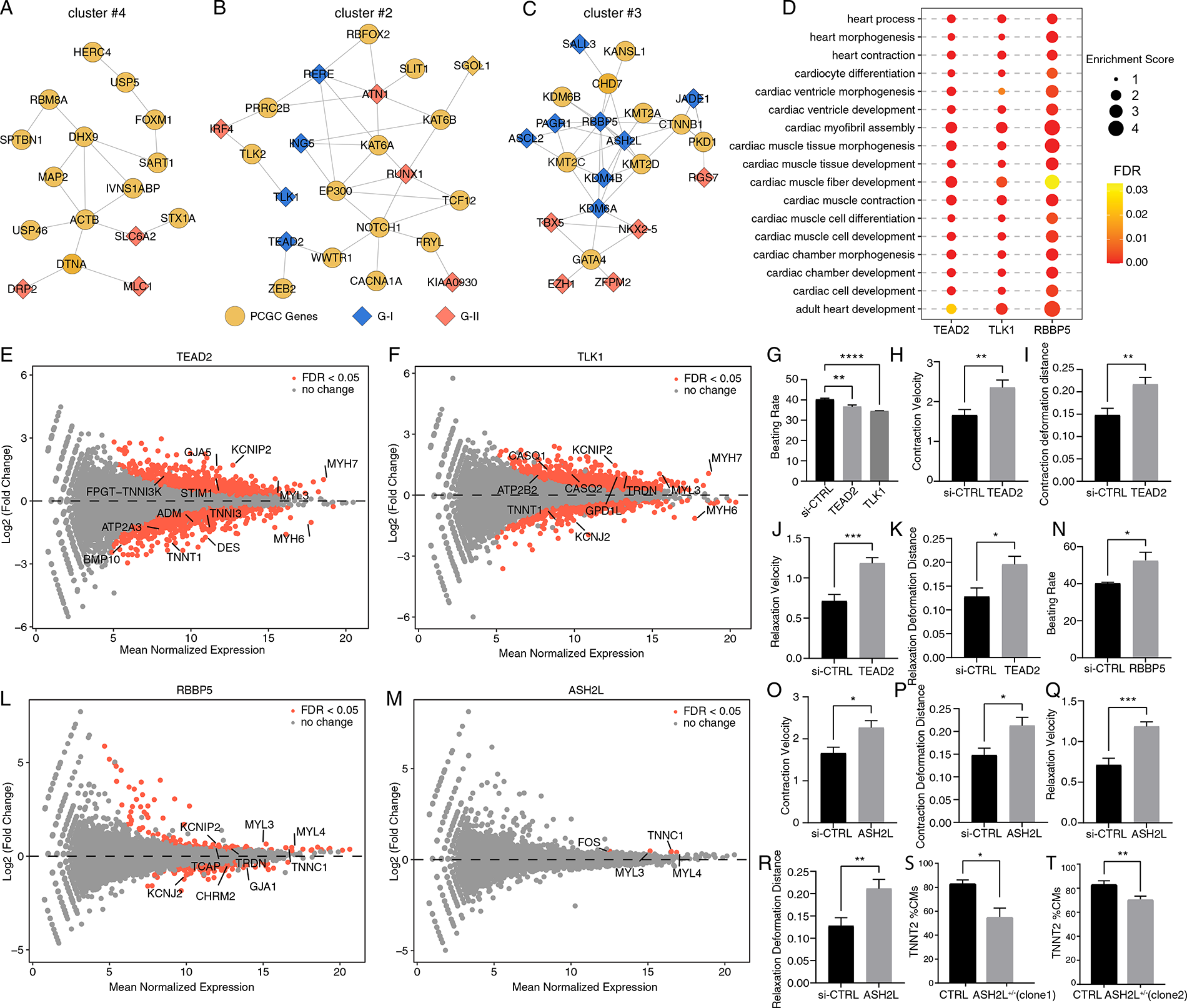
Validating novel functions of the identified genes in regulating fetal heart development. **A-C.** Network clustering identified 33 local clustering structures on the identified network, where cluster #4 **(A)**, #2 **(B)**, #3 **(C)** were presented as representative pathways regulating heart development. **D.** Gene ontology enrichment of the differentially expressed genes in iPSC-CMs upon siRNA knockdown of TEAD2, TLK1 and RBBP5, respectively. These differentially expressed genes consistently displayed strong functional enrichment for heart development and cardiac muscle contraction. The color intensities of the circles represent false discovery rates (FDRs). Sizes of the circles represent the enrichment scores. **E,F,L,M.** RNA-seq identified differentially expressed genes in iPSC-CMs upon siRNA knockdown of TEAD2 **(E),** TLK1 **(F),** RBBP5 **(L)** and ASH2L **(M)**, respectively. X-axis is the mean expression of each gene in iPSC-CMs, and Y-axis indicates their respective fold changes upon siRNA knockdown (siRNA treatment vs. siRNA control). Genes with false discovery rates (FDRs) less than 0.05 were highlighted in red. **G-K.** Cellular contractility assay in the iPSC-CMs. siRNA knockdown against TEAD2 in iPSC-CMs resulted in a marked reduction of the cardiomyocyte beating rate **(G),** increased contraction velocity **(H),** contraction deformation distance(I), relaxation velocity **(J)** and relaxation deformation distance **(K)** relative to the siRNA control. P-values were derived from t-test. **N.** Cellular contractility assay in the iPSC-CMs. RBBP5 knockdown in iPSC-CMs displayed an increased beating rate. P-values were derived from t-test. **O-R**. Cellular contractility assay in the iPSC-CMs. ASH2L knockdown in iPSC-CMs showed increased contraction velocity **(O)**, contraction deformation distance **(P),** relaxation velocity **(Q)** and relaxation deformation distance **(R)**. P-values were derived from t-test. **S-T**. ASH2L^+/-^ knockout lines clone 1 (**S**) and clone 2 (**T**) significantly reduced the differentiation efficiencies into cardiomyocytes (TNNT2 positive cells) from iPSCs. P-values were derived from t-test.

The identified network also encompassed a cluster structure implicating NOTCH1-mediated signaling (cluster #2 in **Table S3**, **Figure 4B**), a master regulator of numerous development processes, including both heart and brain development^62^. NOTCH1 is a known CHD factor and was affected by a de novo LoF mutation in this PCGC cohort. We prioritized to experimentally validate two Group-I genes in this cluster (given clear cardiac functions for Group-II genes, **Table 1**): TEAD2 (TEA Domain Transcription Factor 2) and TLK1 (Tousled Like Kinase 1). TEAD2 is a transcription factor implicated in Hippo signaling, while TLK1 is a kinase regulating chromatin assembly. Notably, TEAD2, as a member of the Group-I gene (**Figure 2B** and **Table 1**), has been characterized as a regulator of neural development^63^, where Tead2^-/-^ mouse mutants displayed exencephaly and open neural tube defects^64^. However, both TEAD2 and TLK1 have uncharacterized function in regulating heart development. We examined their cardiac functions using induced pluripotent stem cell-derived cardiomyocytes (iPSC-CMs). We converted human iPSCs into cardiomyocytes (**Methods and Materials**), and subsequently determined high protein abundance of TEAD2 and TLK1 in the iPSC-CMs (**Figure S2A**). We performed siRNA knockdown to suppress TLK1 and TEAD2 expression, respectively, followed by RNA-seq and cellular assays to determine their regulatory and phenotypic effects in cardiomyocytes. Confirming the knockdown efficiencies by their respective siRNAs (**Figure S3**), we observed that massive genes associated with cardiac muscle contraction exhibited differential expression upon TEAD2 or TLK1 knockdown in cardiomyocytes (**Figure 4D-F** and **Table S4**). Specifically, for TEAD2, despite its known function in regulating neural development, its knockdown in cardiomyocytes has clearly perturbed numerous genes specific for regulating cardiac muscle contractility, including the cardiac muscle myosin factors (MYL3, MYH6/7), and the troponin complex subunits (TNNI3 and TNNT1), the muscle intermediate filament protein (DES) and the connexin gap junction protein (GJA5, **Figure 4E**). Performing cellular contractility assay in the iPSC-CMs, as expected, we observed that the siRNA knockdown against TEAD2 resulted in a marked reduction of the cardiomyocyte beating rate relative to the siRNA control (p = 1.1e-3, **Figure 4G**), accompanied with significantly increased contraction velocity, contraction deformation distance, relaxation velocity and relaxation deformation distance (P < 0.05, **Figure 4H-K**). Similar observation was also made from the TLK1 knockdown experiments, where numerous muscle contractility genes displayed differential expression in the iPSC-CMs (**Figure 4D and F**). In the meantime, knocking down TLK1 expression resulted in reduced beating rate of cardiomyocytes (**Figure 4G**), confirming the role of TLK1 in regulating heart muscle contraction.

The CHD network also encompassed a cluster structure mediated by key factors (GATA4, TBX5, and NKX2-5) regulating heart development (cluster #3 in **Table S3,** **Figure 4C**). However, our analysis now revealed their extensive interactions with numerous histone modification proteins (e.g., KDM6A, KDM4B, ASH2L, RBBP5 and JADE1). We particularly note that many member proteins in this cluster are also implicated in brain development despite three key regulators of heart development GATA4, TBX5, and NKX2-5. For example, GATA4, TBX5, and NKX2-5 all interacted with KDM6A, the causal gene of Kabuki syndrome^65^, characterized by intellectual disability, distinctive facial features and growth delay, etc. Notably, coarctation of the aorta and ventricular/atrial septal defects are also common clinical manifestations of Kabuki syndrome^66^. Indeed, KDM6A mouse mutants also displayed open neural tube defects accompanied with multiple cardiac anomalies including failure of heart looping, thin ventricular wall and myocardial trabeculae hypoplasia (Mouse Genome Informatics^60^). Therefore, this local network structure suggested a potential mechanism of the heart defects in the Kabuki syndrome by perturbing GATA4/TBX5/NKX2-5-mediated cardiac functions through their interactions with KDM6A. More interestingly, this cluster structure also encompassed CHD7, the causal gene for CHARGE syndrome where heart defects and growth retardation are common among patients. This cluster therefore represent a convergent structure underlying several distinct but related monogenic disorders.

To demonstrate the overall implication of this cluster in regulating heart development, we prioritized two Group-I genes, RBBP5 and ASH2L, to experimentally validate their function in heart development given their known function in regulating brain development but uncharacterized cardiac functions. RBBP5 plays a key role in differentiating embryonic stem cells along the neural lineage (UniProtKB/Swiss-Prot Summary), whereas ASH2L regulates the corticogenesis process^67^, neuronal structure and behavior^68^ . Both RBBP5 (RB Binding Protein 5) and ASH2L (ASH2 Like) encode subunits of the histone lysine methyltransferase complex. We performed the same siRNA knockdown experiments and cellular contractility assays as described above. We first confirmed high protein abundance of RBBP5 and ASH2L in iPSC-CMs (**Figure S2B**). Our subsequent RNA-seq revealed differential expression of numerous cardiac muscle proteins in the cardiomyocytes upon RBBP5 knockdown (**Figure 4D and L**), resulting in substantially increased beating rate of the iPSC-CMs (**Figure 4N**). ASH2L knockdown led to fewer differentially expressed genes, but those affected genes were critical factors modulating heart contraction including the myosin light chain proteins MYL3/4, and the troponin subunit TNNC1 implicated in cardiomyopathy^69^. Knocking down ASH2L in iPSC-CMs did not significantly alter the beating rate, but substantially altered all other contractility parameters, including velocity and deformation distance for contraction and relaxation, respectively (**Figure 4O-R**). To gain deeper insights, we further asked whether ASH2L modulates cardiomyocytes differentiation from iPSCs. We generated heterozygous ASH2L knockout iPSC (ASH2L^+/-^) using the CRISPR-Cas9 gene engineering technique. The editing effects were verified by Sanger sequencing and western blot in iPSCs for the two ASH2L^+/-^ knockout clones (**Figure S4**). We then differentiated the edited iPSCs into cardiomyocytes. On day 11, we observed that in both ASH2L^+/-^ knockout lines, the heterozygous ASH2L knockouts indeed significantly reduced the differentiation efficiencies into cardiomyocytes (TNNT2 positive cells) from iPSCs, thereby confirming the role of ASH2L in regulating the cardiogenesis process (**Figure 4S and T**).

Taken together, because Group-II genes were enriched for heart-specific functions (**Table 1**), our study prioritized four Group-I genes (TEAD2, TLK1, RBBP5 and ASH2L) for experimental validation, whose cardiac functions had not been previously characterized. We particularly note RBBP5 and ASH2L, where previous work in mouse suggested Rbbp5 involvement in epigenetic regulation in murine cardiomyocytes^70^ and Ash2l interaction with Tbx1 in vitro^71^. Because RBBP5 is a subunit of histone lysine methyltransferase complex and is widely expressed across human tissues, its role of epigenetic regulation across many cell types (including cardiomyocytes) is anticipated. However, given its well characterized role in differentiating stem cells along the neural lineage, its dual function in regulating heart development was unexpected and is now revealed by our study. For Ash2l interaction with Tbx1^71^, because Tbx1 is a key gene in 22q11.2 deletion syndrome, interacting with Tbx1 likely contribute to the role of Ash2l in regulating heart development. Nevertheless, our experimental data not only demonstrated the effectiveness of our network biology approach on identifying novel genes in regulating fetal heart development, but also provides direct mechanistic insights into the overlapping molecular basis between the brain and heart developmental programs.

### The network is enriched for pathogenic mutations in HLHS probands

Having established the implication of the identified network in regulating heart development, we sought to identify the etiological associations with CHD. Because these genes were not identified from the existing PCGC mutation analysis^10^, we examined the latest release of the PCGC sequencing data (as of Feb, 2021). We performed analyses on patients’ clinical records, and included patients with Tetralogy of Fallot (TOF, N = 328), ventricular septum defects (VSD, N = 776), atrial septum defects (ASD, N = 830), hypoplastic left heart syndrome (HLHS, N = 224), and transposition of the great arteries (TGA, N = 167). Note that all these PCGC proband samples were subjected to whole-exome sequencing on the same platform (Illumina 2000). We retrieved the deposited FASTQ data in PCGC from dbGAP (dbGAP-24034, gap_accession: phs000571, gap_parent_phs: phs001194, SRP025159), and performed variant call and variant annotation (**Methods and Materials**). The variant calls were made by aggregating the whole-exome data for 1,817 unaffected siblings (control subjects, **Methods and Materials**, **Table S5**), which were also used as controls in the previous PCGC study^10^. The joint variant call minimized potential bias from different variant call platforms. We performed additional quality control procedures, which demonstrated high-quality of the identified variants from our variant call procedures **(Figure S5** and **Methods and Materials**).

Using all the called variants, we first confirmed similar population structure between cases and controls (**Figure S6**). Due to weak effect sizes of common variants, we focused our analyses on rare variants from the PCGC cohort. We considered rare variants that were not present in the 1000 Genome database. For each CHD sub-type, we compared deleteriousness of non-synonymous mutations (LoF and missense) in probands and controls. We used the well-known CADD phased scores^32^ to quantify mutational deleteriousness in exonic regions, and computed the mean CADD scores for non-synonymous mutations affecting the network genes in each personal exome in each CHD sub-type (TOF, VSD, ASD, HLHS and TGA) as well as in the control cohort (the unaffected siblings). We reasoned that if the network genes (the orange nodes in **Figure 2A**) were implicated in CHD, we would expect that CHD probands tend to carry more pathogenic mutations affecting the network genes relative to controls. Given differential expression enrichment of the network genes at specific heart development stages (**Figure 2B**), observation of excessive pathogenic mutations specifically affecting particular sub-groups (genes in Group-I or II, **Figure 2B**) among probands of a given CHD sub-type would indicate developmental timing. We separately analyzed Group-I and II genes on the network (**Figure 2B**), and compared the mean CADD scores affecting Group-I or II genes in each proband and control subject. While probands across different CHD subtypes did not display mutational enrichment affecting Group-I genes (**Figure S7**), we did observe significant elevation of mutational pathogenicity affecting Group-II genes, which was specific to HLHS probands (p = 9.7e-3, Wilcoxon rank-sum test, **Figure 5A**) but was insignificant for other CHD subtypes in comparison (**Figure 5A**). Because HLHS is typically comorbid with ASD, we indeed observed that 56 HLHS probands (among total 224 HLHS cases) also received ASD diagnosis. Excluding these overlapping cases further boosted the statistical significance for the enrichment of pathogenic non-synonymous mutations in the identified network genes (p = 2.7e-3, Wilcoxon rank-sum test, **Figure 5C**). We performed a set of additional control experiments to confirm the observation: (1) we performed the same analysis on rare synonymous mutations affecting Group-II genes, but did not observe any statistical significance among all CHD sub-types, including HLHS (**Figure 5B**). This observation demonstrated that the observed statistical significance indeed implied functional consequences. (2) We performed the same analysis on 62 lung-specific protein-coding genes^34^, and no significance was observed on both non-synonymous (p = 0.86, Wilcoxon rank-sum test) and synonymous (p = 0.36, Wilcoxon rank-sum test) variants (**Figure 5D**). This comparison confirmed specificity of the identified genes in regulating heart development. (3) In each exome, we confirmed that the number of rare non-synonymous variants in each HLHS proband did not significantly differ from the unaffected siblings in the control cohort (p = 0.55, Wilcoxon rank-sum test, **Figure S8**), suggesting that the observed enrichment of pathogenic mutations cannot be explained by excessive mutations identified in the proband cohort than the control cohort, but by the increased mutational pathogenicity. (4) Lastly, we asked whether our observation could be merely explained by increased CADD scores of all rare non-synonymous variants across the exome background in cases relative to controls. We performed a permutation study, where, in each permutation, we randomly sampled rare non-synonymous variants from the exome backgrounds from cases and control cohorts, matching the number of rare non-synonymous variants in Group-II genes in cases and control cohorts, respectively. We performed the permutation 100 times, and confirmed that mutational pathogenicity scores were not significantly different between cases and controls when randomly sampling rare non-synonymous variants from the two cohorts (**Figure 5E**). Taken together, our comparisons collectively demonstrated the specificity of the identified network in the molecular etiologies of HLHS.

**Figure 5.**
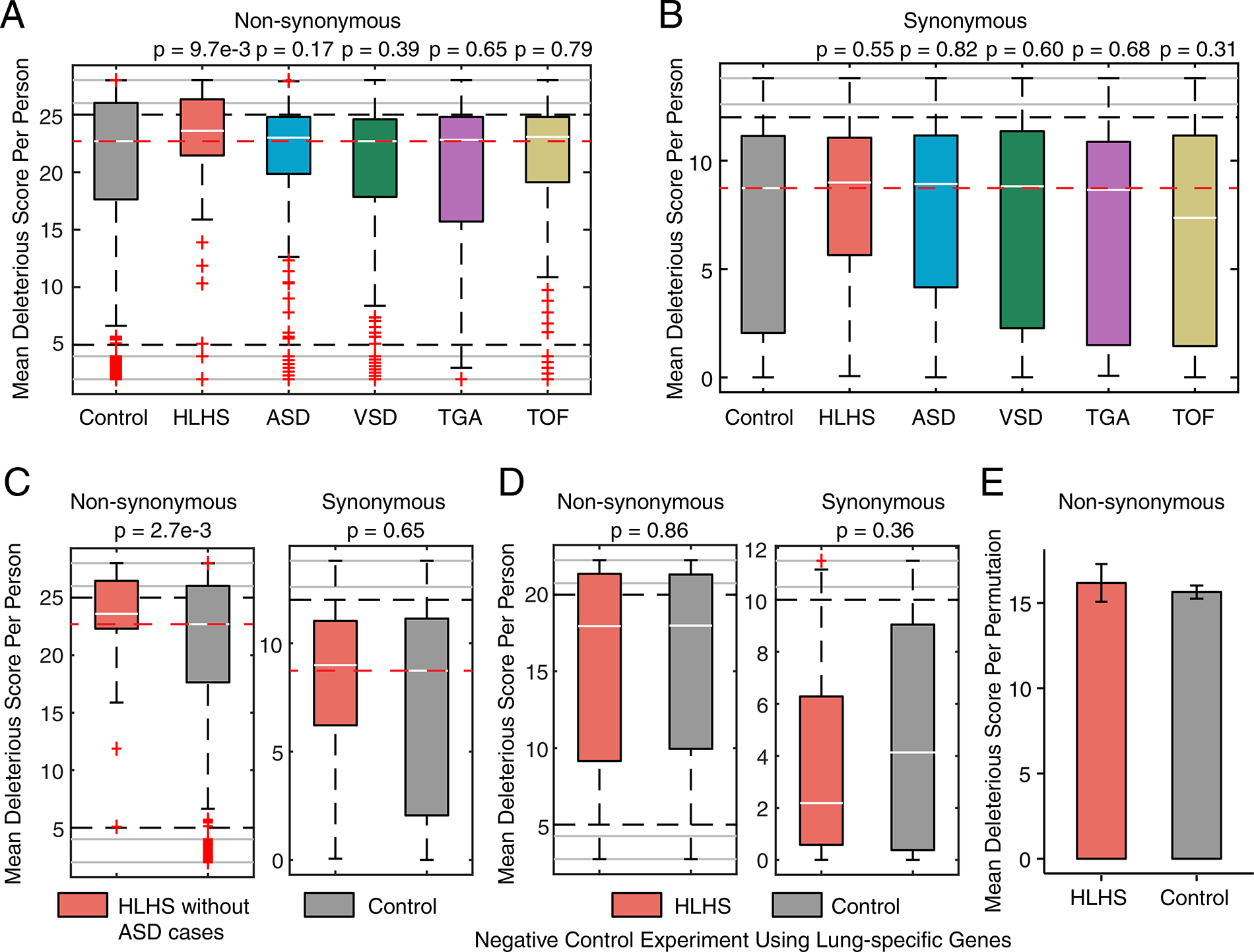
Group-II genes on the network were enriched for pathogenic mutations in probands with HLHS. **A.** Rare non-synonymous variants displayed a significant increase in mutational deleteriousness in HLHS probands, but not in individuals with ASD, VSD, TGA or TOF, relative to unaffected siblings. **B.** Significant differences in mutational deleteriousness of rare synonymous mutations were not observed in any CHD subtypes relative to unaffected siblings. HLHS, ASD, VSD, TGA and TOF stand for hypoplastic left heart syndrome, atrial septal defects, ventricular septal defects, transposition of the great arteries, and tetralogy of fallot. Mutational pathogenicity was measured by CADD scores. We computed the mean CADD scores for non-synonymous in Group-II genes in each personal exome, and we compared the mean CADD score distribution among probands in each CHD subtype against the distribution among the unaffected siblings. The same comparison was performed on synonymous variants as a set of negative controls. Controls were the unaffected siblings from the original publication^10^. P-values were derived from Wilcoxon rank-sum test. **C.** Excluding individuals comorbid with ASD further boosted statistical significance (Group-II genes). **D.** The statistical significance was absent on lung-specific genes for both non-synonymous and synonymous variants. **E.** Permutation analysis confirmed that CADD scores had a similar distribution for rare non-synonymous variants in the background exomes in the HLHS cohort relative to the control cohort. In each permutation, rare non-synonymous variants were randomly sampled from the exome backgrounds in the HLHS and control cohorts respectively, matching the number of rare non-synonymous variants in Group-II genes in cases or in controls, followed a comparison between their CADD scores using Wilcoxon rank-sum test. Among 100 permutations, 98 were statistically insignificant (p = 0.98), suggesting that our observation cannot be merely explained by exome background differences in CADD scores. Error bars represent standard error of the mean.

### Single cell analysis of the CHD network in HLHS

Since HLHS is a form of critical congenital heart defect (CCHD), we next sought to derive further mechanistic insights into the molecular etiologies of HLHS. HLHS affects blood flow through the heart due to the underdeveloped left ventricle accompanied with malformations of mitral and aortic valves^72^. To understand the association of the network with HLHS, we leveraged the recently published single-cell data from an HLHS fetal heart (day 84), and compared gene expression in the underdeveloped left ventricle of this HLHS heart against the left ventricle of a typically developing fetal heart on day 83 (**Methods and Materials**)^35^. For Group-I and II genes in the network (**Figure 2A**), we compared their expression in the HLHS heart across all cell types. We did not observe significant expression differences in the HLHS heart for Group-I genes across all cell types, which was expected given their expression specificity in early developmental stages (compared with the HLHS heart from day 84). However, for Group-II genes, we observed their significant expression reduction only in endothelium cells of the HLHS left ventricle across all cell types (p = 5.8e-4, Wilcoxson rank-sum test, **Figure 6A**). Close examining the Group-II genes, we observed that the reduction was consistent between the two sub-groups (Group-II-A and Group-II-B, **Figure 2B**), but was particularly pronounced among Group-II-A genes (**Figure 6B**) whose expression were specific across fetal development stages (**Figure 3**), thereby highlighting the significant contribution to HLHS from fetal endothelium development. We also noted the conduction system in **Figure 6A**, where expression of Group-II genes displayed marginal statistical significance (p = 0.08, Wilcoxon rank-sum test). Close examination revealed a significant expression reduction only specific to Group-II-A genes in the conduction system in the HLHS heart (left ventricle) (**Figure 6C**), again suggesting a dysregulation of the conduction system in HLHS. Taken together, our analysis thus strongly suggests the impaired endothelium and conduction system in HLHS.

**Figure 6.**
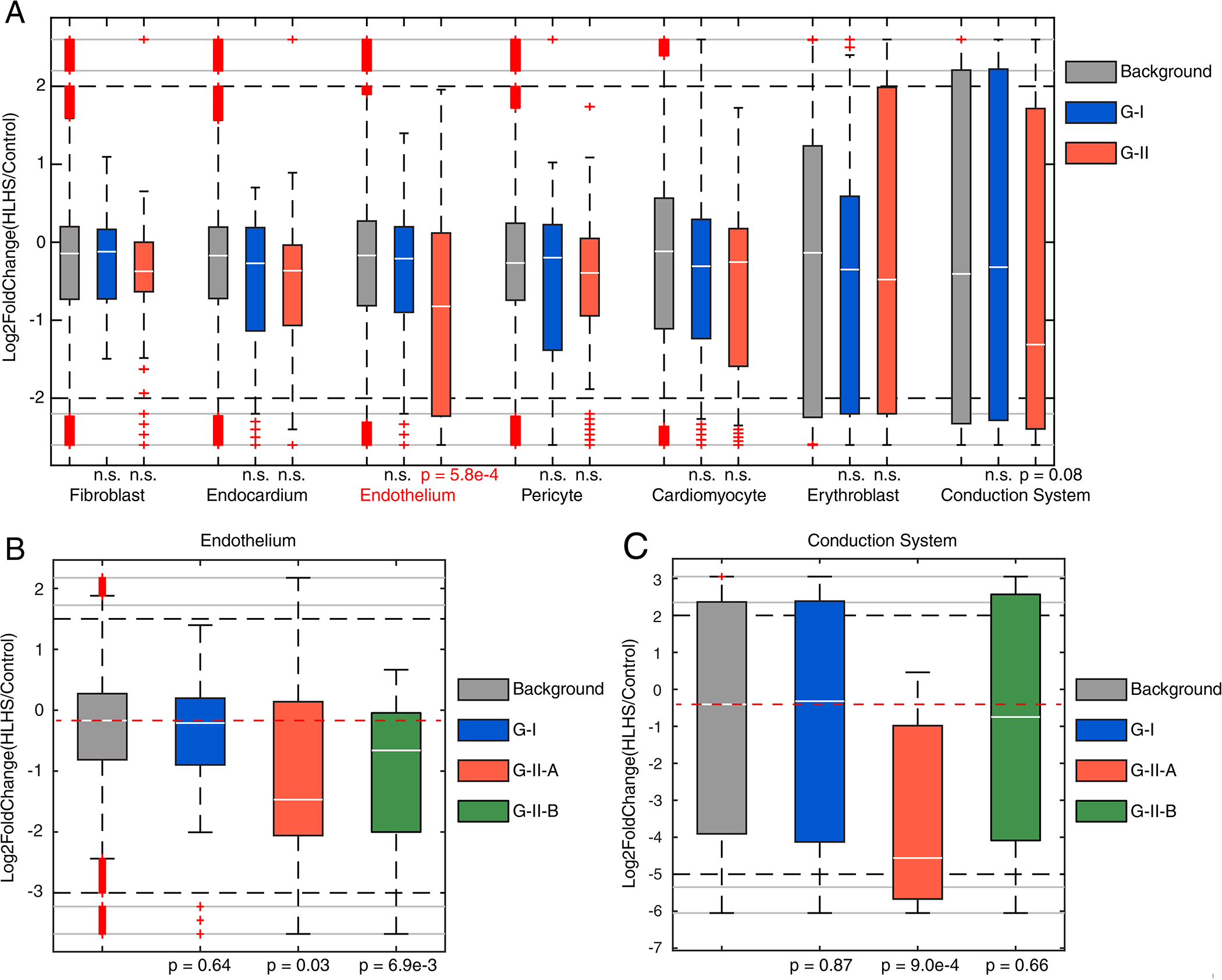
Single-cell analysis of the network in the HLHS heart. **A.** Across all the cell types in the HLHS left ventricle, Group-II (G-II) genes showed the strongest expression reduction in the endothelium cells. P-values were derived from Wilcoxon rank-sum test, **B.** Examining two subgroups of Group-II (G-II) genes revealed significant downregulation of both subgroups (G-II-A and G-II-B) in the endothelium cells. **C.** Only Group-II-A (G-II-A) genes displayed significant expression reduction in the conduction system in the HLHS left ventricle.

## Discussion

Disease-associated mutations are not randomly dispersed across the genome but affect common sets of molecular pathways on biological networks leading to common clinical manifestations^12^. The systems thinking has motivated us to develop a novel computational framework to integrate large-scale CHD genomes, the human interactome, the fetal heart spatial transcriptome as well as the single-cell transcriptome from clinical samples. This integrative strategy identified numerous novel proteins with previously uncharacterized roles in regulating fetal heart development, and our subsequent multi-omic analyses further demonstrated their function more specific to certain CHD subtypes with the strongest effect on HLHS. Overall, our integrative analysis significantly advanced our understanding of the genetic architecture in CHD, revealed the molecular etiologies in HLHS, and can be readily extended to study other complex diseases.

We started our analysis by seeding CHD candidate proteins on the human protein interaction network. These proteins were intolerant to copy losses and were affected by de novo LoF mutations in CHD probands. Their central positions and the clustering patterns on the network suggested their significant impacts and convergent functions in CHD, which has enabled us to implement the NetWalker approach to identify a connected network underlying fetal heart development. We observed two expression clusters (Group-I and II genes, **Figure 2A**) in the identified CHD network. While Group-II genes displayed strong tissue specificity in the fetal heart, it was interesting that Group-I genes, showing the strongest expression at very early developmental stages (**Figure 2A**), modulate both neurodevelopmental and heart developmental programs. This observation was concordant with previous work, where autism genes were also frequently identified as CHD candidate genes^10^; however, in this work, the shared molecular etiologies were only limited to Group-I genes, further demonstrating the functional convergence was specific to early developmental stages. Leveraging the global protein interaction network, we were able to further pinpoint the underlying mechanisms in context of the local interacting proteins. The FOXM1 cluster (**Figure 4A**) best demonstrated the notion of mutational convergence, where the majority of its member proteins were affected by de novo LoF mutations in CHD probands. In the same vein, our experimental validation in iPSC-CMs further confirmed four additional Group-I genes for their previously uncharacterized function in heart development, including the previously characterized factor in neural tube defects, TEAD2, and ASH2L regulating corticogenesis. Our RNA-seq experiments and cellular contractility assays consistently revealed their function in regulating cardiomyocyte contraction, demonstrating novel heart-specific functions of these genes previously recognized as brain genes. In addition to studying mature cardiomyocytes, we also performed gene editing to perturb ASH2L expression in iPSCs and confirmed ASH2L function in controlling the differentiation process into cardiomyocytes. This observation is consistent with SALL3 function in the GATA4 cluster (**Figure 4C**), where SALL3 acted as a switch controlling the developmental trajectory of stem cells towards the neural or cardiac lineage^73^. Taken together, Group-I genes constitute the shared molecular etiologies between cardiac and neurological conditions given their tissue-context-dependent functions or their role in controlling cell differentiation processes.

We analyzed rare mutations from the PCGC cohort on the network, and observed that Group-II genes were specifically enriched for pathogenic mutations from individuals with HLHS. This observation was consistent with our transcriptome analysis of the typically developing and HLHS heart, thereby revealing potential tissue of origins underlying HLHS. We particularly highlight the role of endothelium cells in HLHS, where its etiological contribution has just recently begun to be recognized^35^. Although we demonstrated increased mutational load in Group-II genes in HLHS, this observation did not fully preclude the role of Group-I genes in HLHS development. In fact, among the four Group-I genes (RBBP5, ASHL2, TLK1 and TEAD2) we experimentally validated in iPSC-CMs (**Figure 4**), RBBP5 also displayed significant down-regulation in the cardiomyocytes of this HLHS sample. Because Group-I genes are also associated with neurodevelopmental phenotypes, severe phenotypic consequences likely have suppressed excessive deleterious mutations in the human population, or they likely underlie many syndromic cases (eg. Kabuki syndrome, **Figure 4C**). Therefore, screening individual pathogenic mutations in Group-I genes would provide a molecular basis for clinical evaluation of the comorbidities between CHD and neurodevelopmental conditions.

## Acknowledgment

JL acknowledges the startup fund from the Eli and Edythe Broad Center of Regeneration Medicine and Stem Cell Research, the Bakar Computational Health Sciences Institute, and the Parker Institute for Cancer Immunotherapy at UCSF. MS acknowledges grant award NIH S10OD025212, and NIH/NIDDK P30DK116074.

