## Supplemental Files for "Systems Analysis of de novo Mutations in Congenital Heart Diseases Identified a Molecular Network in Hypoplastic Left Heart Syndrome"

**Supplementary material**

**Table S1. Functional enrichment of PCGC CHD candidate genes**

A. List of 120 PCGC CHD probands genes.

B. Functional enrichment of 120 PCGC CHD probands genes.

C. List of 35 PCGC CHD siblings genes.

D. Functional enrichment of 35 PCGC CHD siblings genes.

**Table S2. Human protein interactions**

**Table S3. 33 new identified network clusters**

**Table S4. Go enrichment of differentially expressed genes in iPSC-CMs upon siRNA knockdown**

A. GO enrichment of si-TEAD2 differential genes.

B. GO enrichment of si-TLK1 differential genes.

C. GO enrichment of si-RBBP5 differential genes.

**Table S5. 1,817 unaffected siblings**

**Table S6. 1,865 PCGC cases information**

B-F columns: 1 indicates the case is included in the corresponding CHD subtype and 0 indicates exclusion.

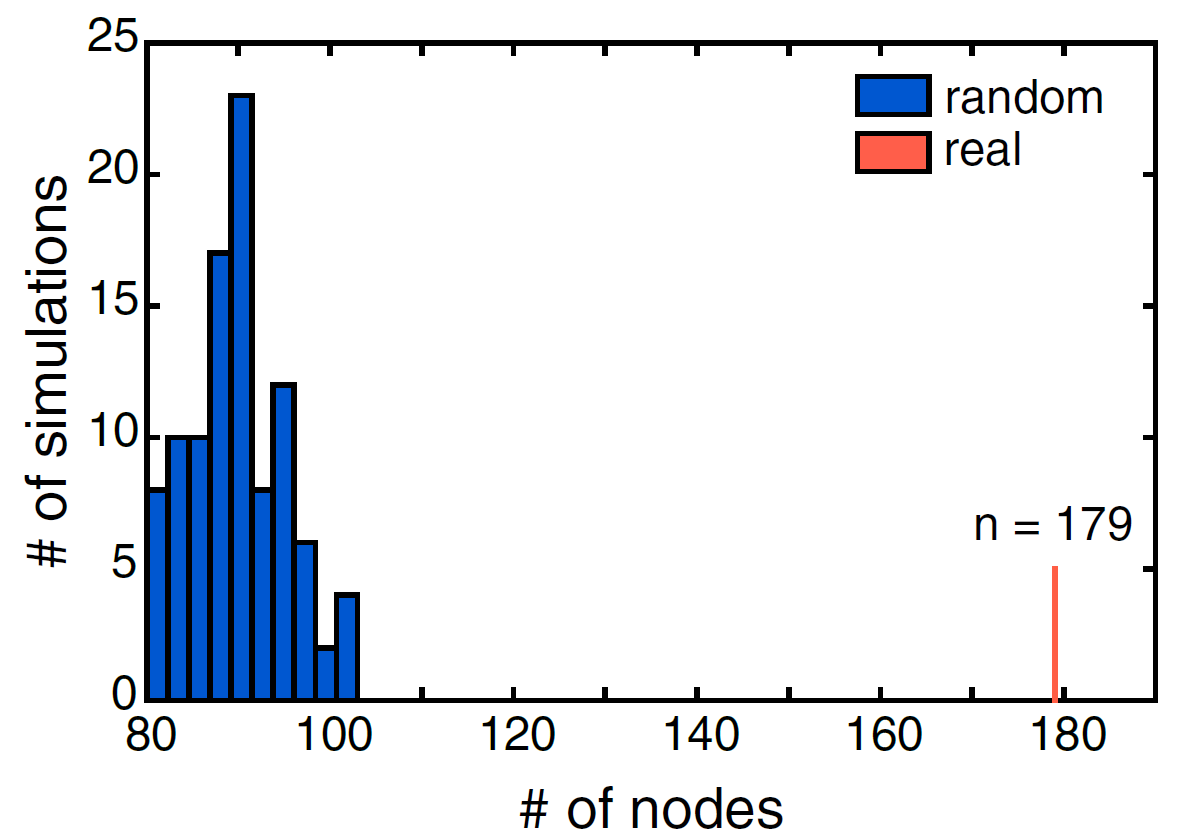

**Figure S1. Distribution of the number of interacting partners on the interaction network.**

100 random permutations of interacting partners for each protein on the interaction network with the same degrees demonstrated the newly identified proteins formed a highly connected network. Numbers of the interacting partners on permutated networks were colored in bule, and the number of interacting partners on the newly identified network were colored in red.

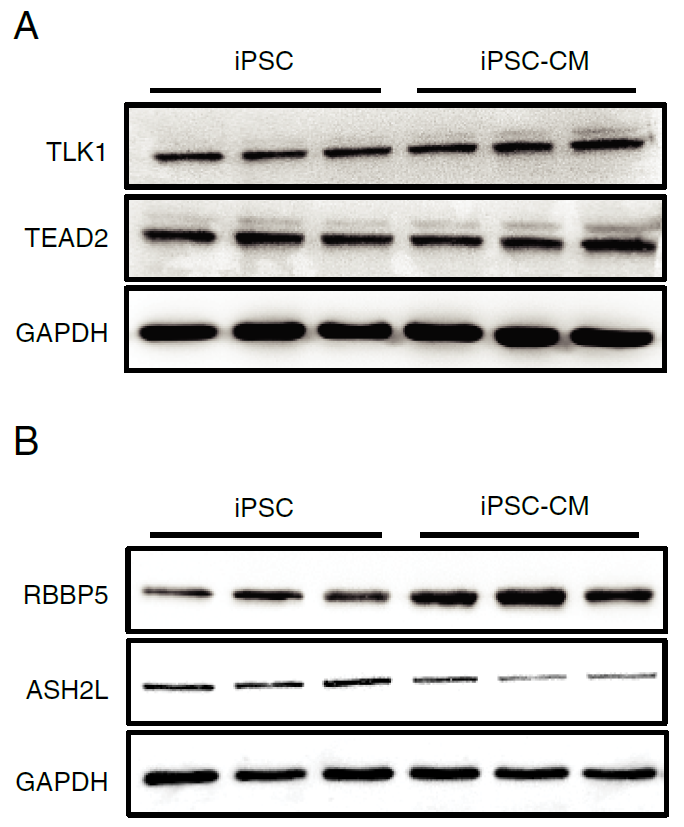

**Figure S2. TLK1, TEAD2, RBBP5 and ASH2L protein expressions.**

**(A-B)** Western blotting was used to determine the proteins expressions. **(A)** Immunoblots of TLK1 (86 kDa), TEAD2 (50 kDa), and GAPDH (36 kDa) in iPSCs (n = 3) and iPSC-CMs (n = 3). These 3 groups of protein blots were cropped from different parts of the same gel. **(B)** Immunoblots of RBBP5 (60 kDa), ASH2L (69 kDa), and GAPDH (36 kDa) in iPSCs (n = 3) and iPSC-CMs (n = 3). These 3 groups of protein blots were cropped from different parts of the same gel.

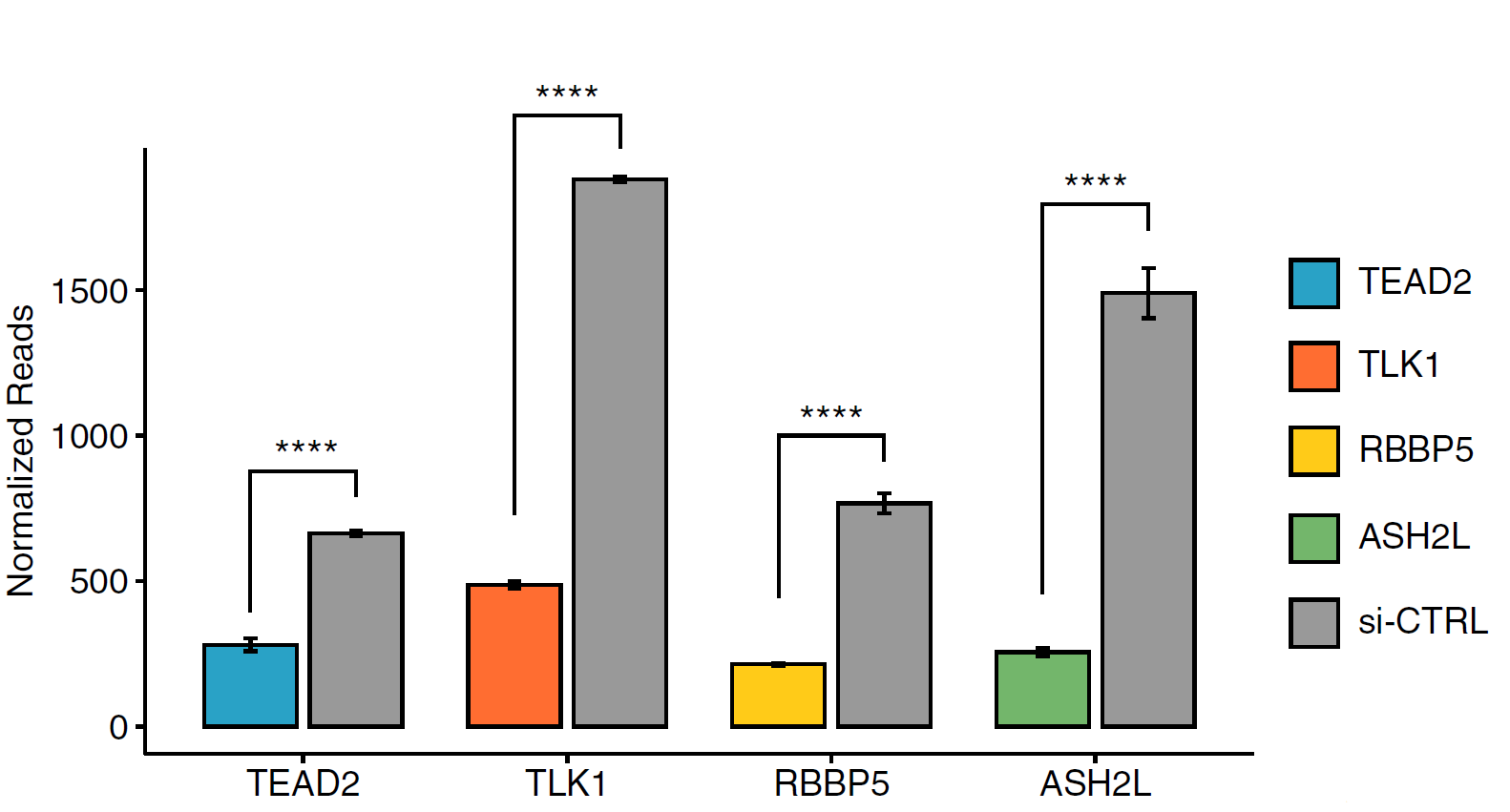

**Figure S3. siRNA knockdown efficiency.**

siRNA knockdown of TEAD2, TLK1, RBBP5 and ASH2L showed efficient reduction of the target mRNA level, respectively. P-values were derived from t-test. Error bars represent standard error of the mean.

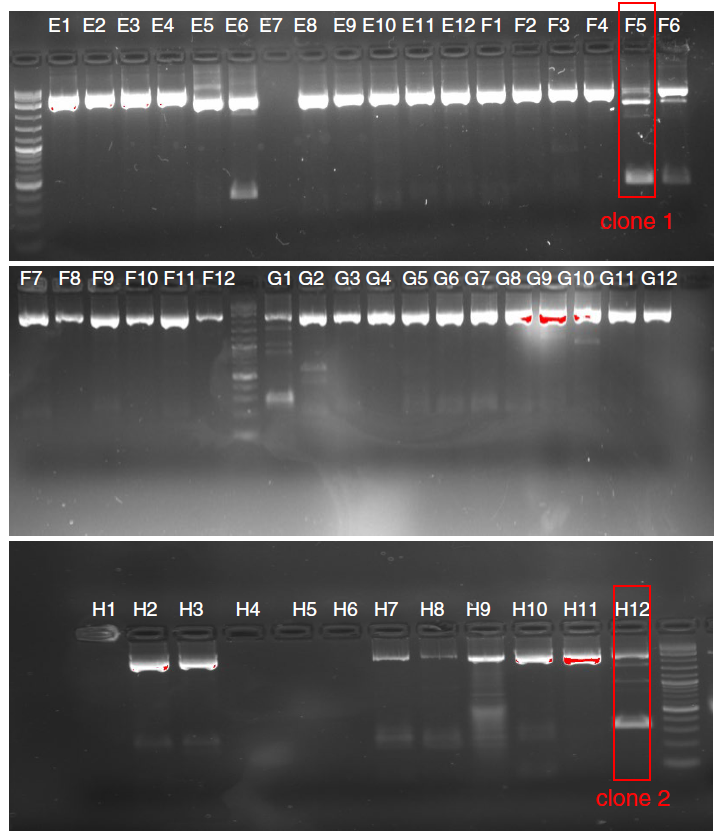

**Figure S4. ASH2L+/- knockout clones editing effects.**

Two ASH2L^+/-^ knockout clones (clone1 and clone2) editing effects were verified by western blot in iPSCs.

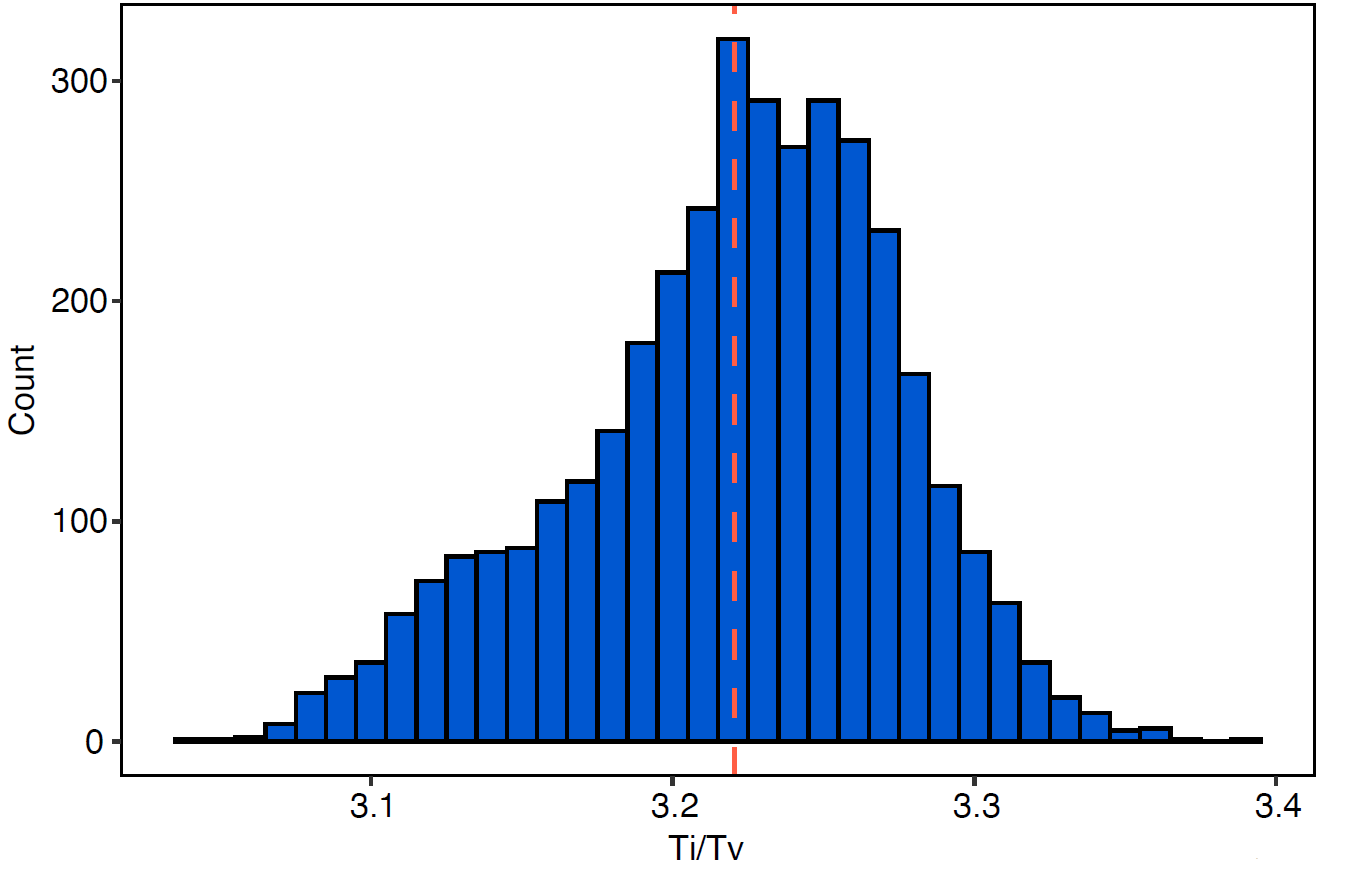

**Figure S5. Quality control of the called variants.**

The identified variants from our variant call procedures showed high quality. The distribution of Ti/Tv ratios across 1,865 cases and 1,817 control subjects with a mean of 3.22.

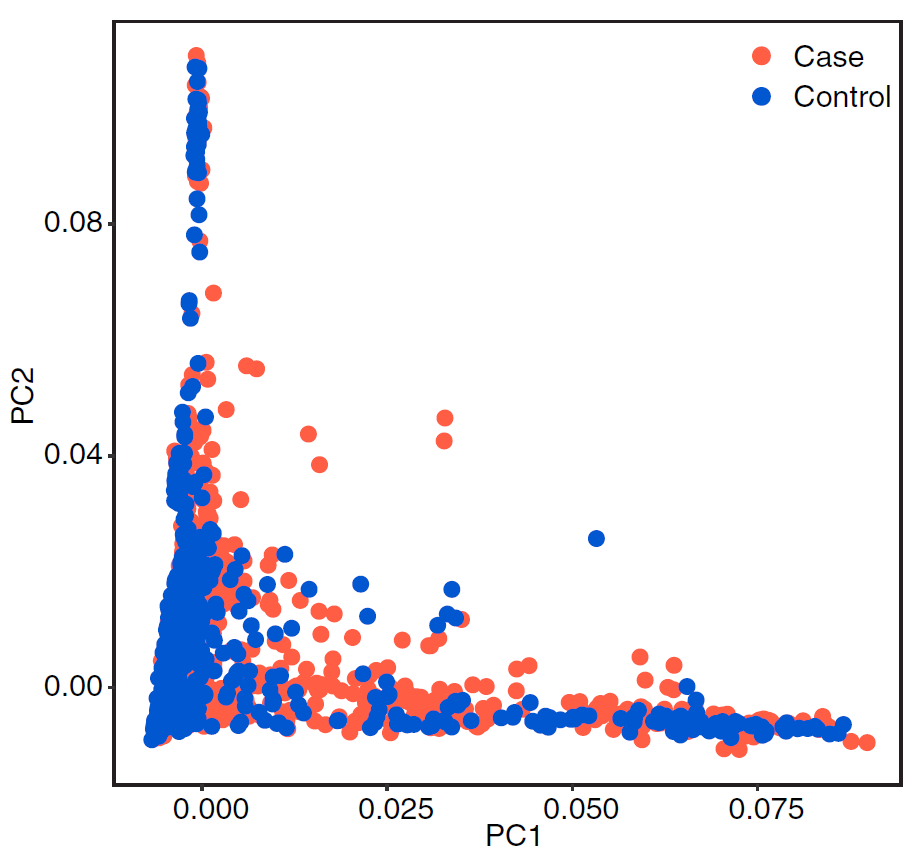

**Figure S6. Population structure Analysis.**

PCA analysis on all the called exonic variants revealed similar population structure between cases (n = 1,865, red) and control subjects (N = 1,817, blue).

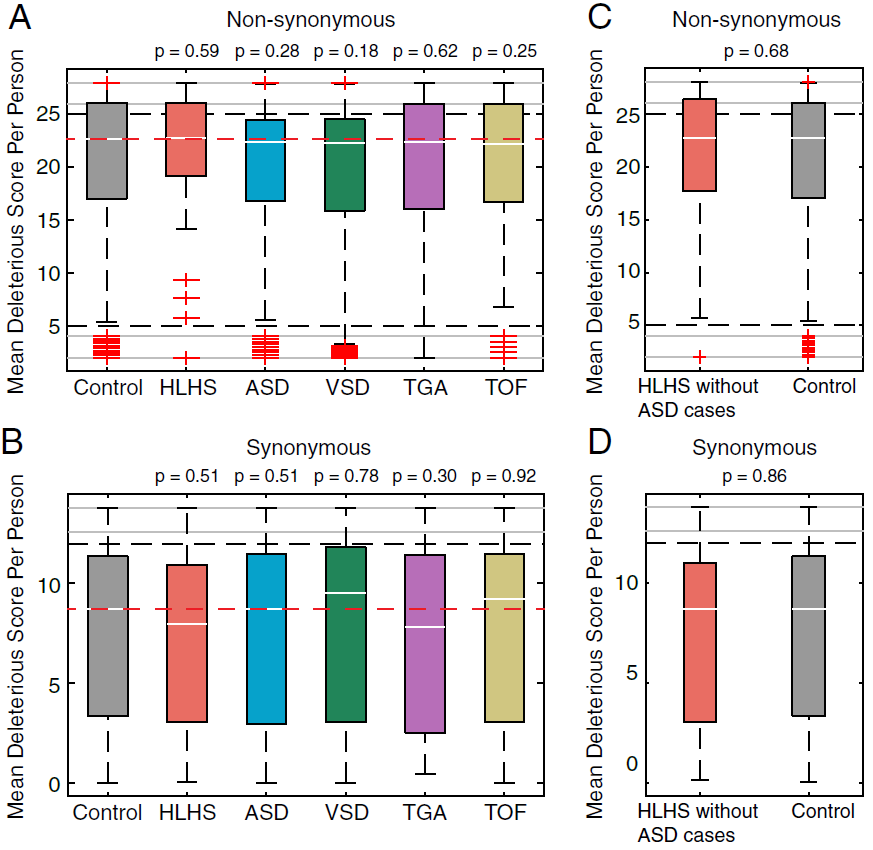

**Figure S7. Group-I genes on the network did not display increased mutational deleteriousness in CHD probands relative to unaffected siblings.** The comparisons were performed on rare non-synonymous mutations **(A)** and rare synonymous mutations **(B).** HLHS, ASD, VSD, TGA and TOF stand for hypoplastic left heart syndrome, atrial septal defects, ventricular septal defects, transposition of the great arteries, and tetralogy of fallot. Mutational pathogenicity was measured by CADD scores. We computed the mean CADD scores for non-synonymous **(A)** and synonymous **(B)** in Group-I genes in each personal exome, and we compared the mean CADD score distribution among probands in each CHD subtype against the distribution among the unaffected siblings. The same analysis was also performed on HLHS cases after excluding those comorbid with ASD, but no significance was observed on rare non-synonymous (**C**) and synonymous variants (**D**). P-values were derived from Wilcoxon rank-sum test.

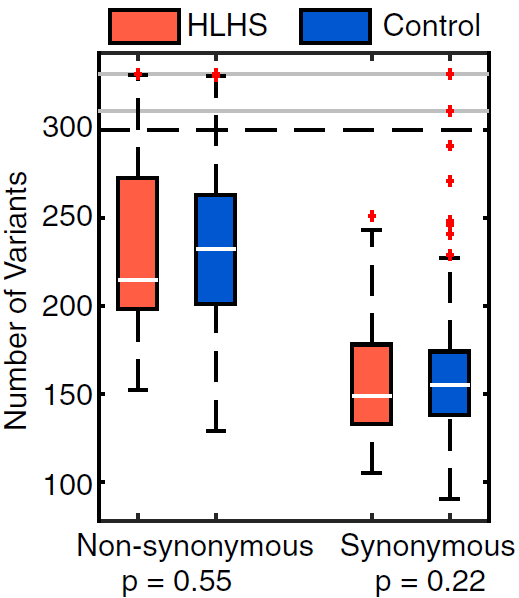

**Figure S8. The number of rare variants in exome did not show significant differences between HLHS probands and control cohorts.**

The number of non-synonymous and synonymous variants in each HLHS proband did not significantly differ from the unaffected siblings in the control cohort. HLHS stand for hypoplastic left heart syndrome. P-values were derived from Wilcoxon rank-sum test.
